# Evolution of extended-spectrum β-lactamase-producing ST131 *Escherichia coli* at a single hospital over 15 years

**DOI:** 10.1101/2023.12.11.571174

**Authors:** Shu-Ting Cho, Emma G. Mills, Marissa P. Griffith, Hayley R. Nordstrom, Christi L. McElheny, Lee H. Harrison, Yohei Doi, Daria Van Tyne

## Abstract

*Escherichia coli* belonging to sequence type ST131 constitute a globally distributed pandemic lineage that causes multidrug-resistant extra-intestinal infections. ST131 *E. coli* frequently produce extended-spectrum β-lactamases (ESBLs), which confer resistance to many β-lactam antibiotics and make infections difficult to treat. We sequenced the genomes of 154 ESBL-producing *E. coli* clinical isolates belonging to the ST131 lineage from patients at the University of Pittsburgh Medical Center (UPMC) between 2004 and 2018. Isolates belonged to the well described ST131 clades A (8%), B (3%), C1 (33%), and C2 (54%). An additional four isolates belonged to another distinct subclade within clade C and encoded genomic characteristics that have not been previously described. Time-dated phylogenetic analysis estimated that the most recent common ancestor (MRCA) for all clade C isolates from UPMC emerged around 1989, consistent with previous studies. We identified multiple genes potentially under selection in clade C, including the cell wall assembly gene *ftsI*, the LPS biosynthesis gene *arnC*, and the yersiniabactin uptake receptor *fyuA.* Diverse ESBL genes belonging to the *bla*_CTX-M_, *bla*_SHV_, and *bla*_TEM_ families were identified; these genes were found at varying numbers of loci and in variable numbers of copies across isolates. Analysis of ESBL flanking regions revealed diverse mobile elements that varied by ESBL type. Overall, our findings show that ST131 subclades C1 and C2 dominated and were stably maintained among patients in the same hospital and uncover possible signals of ongoing adaptation within the clade C ST131 lineage.

## Introduction

*Escherichia coli* sequence type (ST) 131 is a globally distributed extra-intestinal pathogenic *E. coli* (ExPEC) lineage that causes bloodstream and urinary tract infections^1^. ST131 isolates commonly exhibit multidrug resistance and often produce extended-spectrum β-lactamases (ESBLs), which give them the ability to resist therapy with many β-lactam antibiotics including expanded-spectrum cephalosporins^2^. The emergence and global spread of ESBL-producing *E. coli* raise serious issues for clinical management.

Prior studies have shown that the *E. coli* ST131 population can be separated into three major phylogenetic clades^3^. Typing of the *fimH* locus has been traditionally used to classify isolates into clade A (*fimH*41), clade B (*fimH*22), and clade C (*fimH*30). Isolates belonging to clade A have been mostly found in Asia, whereas clade C isolates dominate in the United States^4^. The clade C population has further diverged into the nested subclades C1 (*fimH*30R) and C2 (*fimH*30Rx), with isolates in both subclades encoding mutations in the *gyrA* and *parC* genes that confer resistance to fluoroquinolones. Most isolates in the C2 subclade carry the ESBL gene *bla*_CTX-M-15_, while isolates in the C1 subclade often carry *bla*_CTX-M-27_^5^. ESBL genes are frequently maintained on mobile genetic elements (MGEs)^6^, which are often carried on plasmids but can also be integrated into the chromosome^7^.

Here we survey the genomic diversity and evolution of ESBL-producing ST131 *E. coli* isolates at a single medical center in the Pittsburgh area over a 15-year period. We describe the distribution of subclades and the diversity of ESBL-encoding MGEs, as well as the evolution of clade C isolates specifically, at our hospital. Our results suggest that a diverse ST131 *E. coli* population circulates in our facility, from which we periodically sampled. We also found evidence that distinct ST131 subpopulations have persisted in our hospital for over a decade, suggesting that multiple subclades are stably maintained in this setting.

## Results

### The ESBL-producing *E. coli* ST131 population at UPMC is dominated by clade C

To survey the genomic diversity of ESBL-producing ST131 *E. coli* at the University of Pittsburgh Medical Center (UPMC), we sequenced the genomes of 154 clinical isolates collected from patients between 2004 and 2018 (Table S1). ESBL-producing *E. coli* isolates collected between 2004 and 2016 were tested with PCR using ST131-specific primers^8^, and up to ten ST131 isolates from each year were selected for whole genome sequencing. Beginning in 2016, isolates were identified as ST131 through analysis of whole genome sequence data generated previously^9^. We included isolates belonging to ST131 based on multi-locus sequence typing (MLST), as well as three isolates that belonged to ST8347 (a single locus variant of ST131) and two isolates that belonged to two additional single locus variants of ST131 that have not yet been assigned a sequence type (Fig. 1).

**Figure 1.**
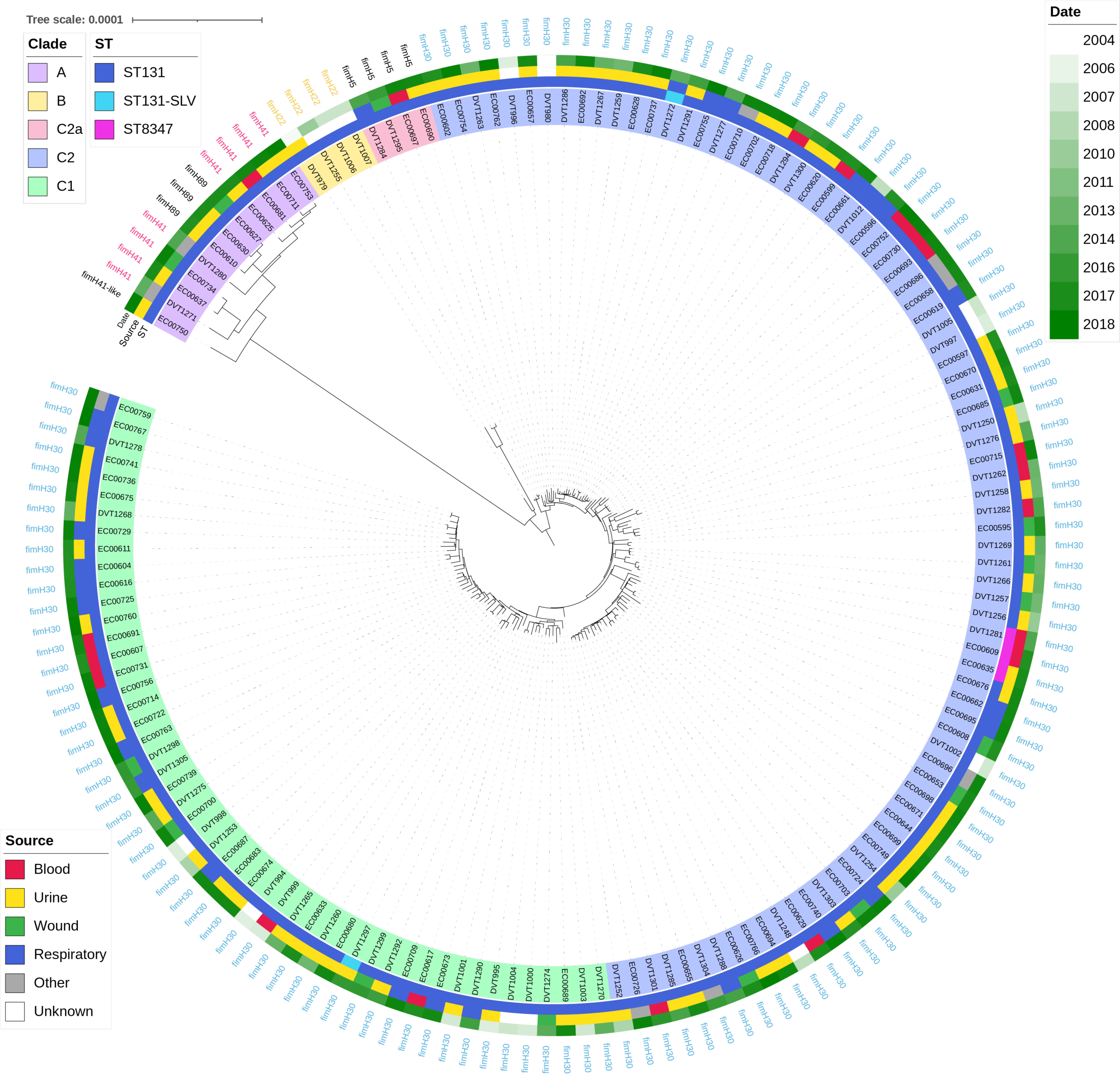
Genetic diversity of 154 ESBL-producing ST131 *E. coli* isolates. The maximum likelihood phylogeny was constructed with RAxML from 18,734 core genome single nucleotide polymorphisms (SNPs). Background shading of each isolate indicates the ST131 clade (A, B), subclade (C2, C2), or subgroup (C2a). Multi-locus sequence type (ST), source, and date of isolation are shown as color blocks next to each isolate. *fimH* alleles were predicted from genome sequences.

A recombination-filtered phylogenetic tree based on variants found in the core genome of all 154 isolates was constructed using RAxML (Fig. 1). As expected for the ST131 population^4,6,10^, isolates resided on three major branches. The first branch (clade A) contained twelve isolates (7.8%), including eight with *fimH*41, three with *fimH*89, and one with a novel *fimH* sequence that was most similar to *fimH*41 (Fig. 1). These isolates were all collected in 2013 and later (Fig. S1). An additional four isolates (2.6%) collected in 2005, 2007, and 2010 encoded *fimH*22 and belonged to clade B. The third branch consisted of the remaining 138 isolates (89.6%), including one group of four isolates that encoded *fimH*5. The rest of the isolates on this branch encoded *fimH*30, indicating that the clade should be assigned as clade C (Fig. 1). QRDR mutations in *gyrA* and *parC* were detected in all 138 clade C isolates. The 86 isolates carrying two mutations described previously^4^ were assigned to subclade C2. Within this clade, the four isolates encoding *fimH*5 were designated as subgroup C2a. The remaining 52 clade C isolates were classified as subclade C1. Clade C isolates were collected throughout the study period and there was no apparent difference in collection dates of subclade C1 versus C2 isolates (Fig. S1).

### Evolution of clade C and stable maintenance of subclades C1 and C2 in the Pittsburgh area

Prior studies have suggested that clade C emerged in approximately 1990^6,10,11^. To examine the evolution of clade C in our hospital, we performed a time-calibrated phylogenetic analysis using TreeTime (Fig. 2)^12^. The estimated substitution rate was 1.76 core genome mutations per genome per year, and the estimated root date of clade C was 1988.5. In addition, when we re-rooted the phylogenetic tree to separate subclades C1 and C2, we confirmed that the C2a subgroup was embedded within subclade C2. The estimated date of emergence of this subgroup from the subclade C2 population was approximately 2013 (Fig. 2).

**Figure 2.**
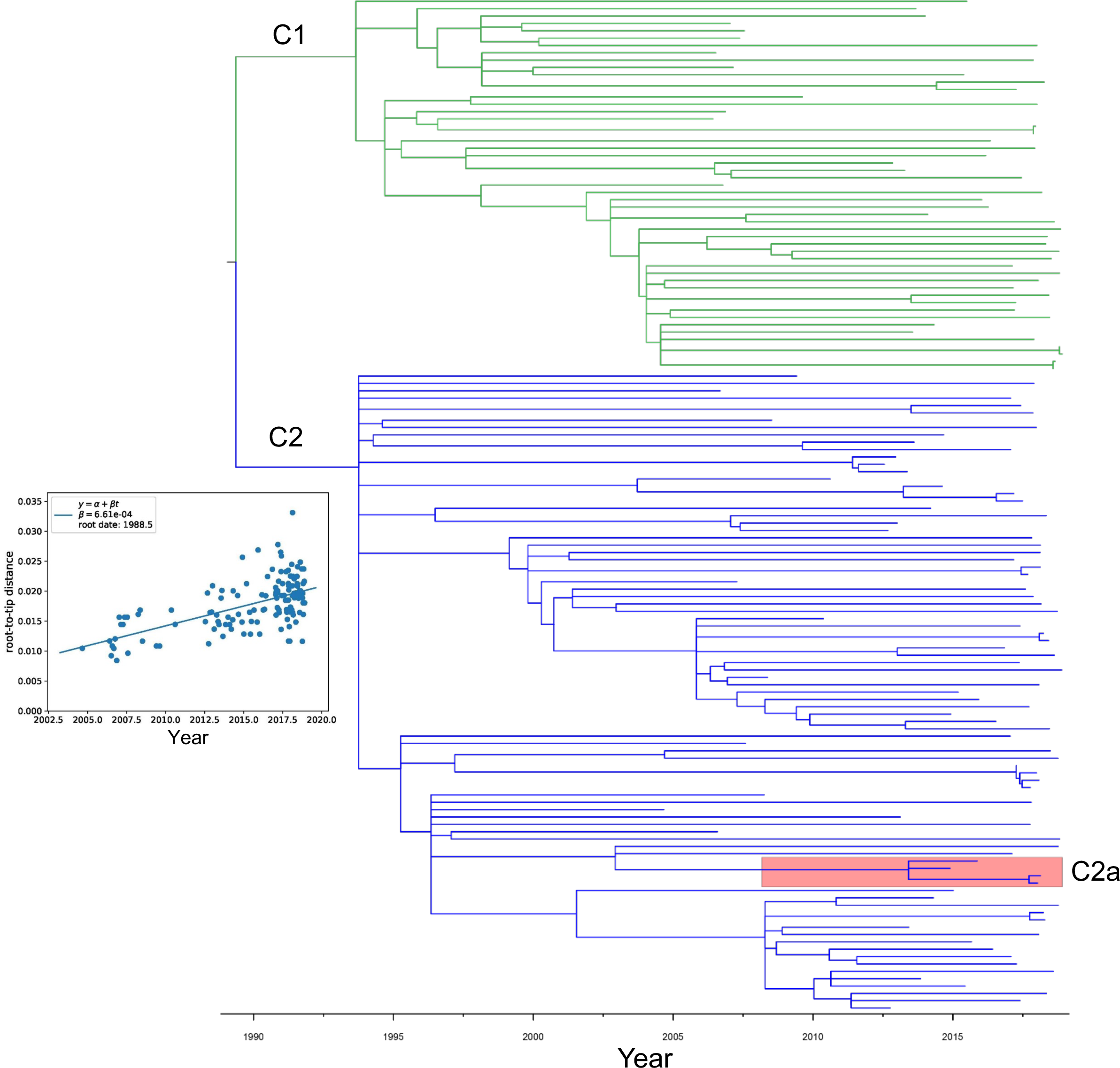
Time-calibrated phylogeny of 138 clade C isolates. The molecular-clock phylogeny was inferred from 2,656 aligned SNPs and was constructed with TreeTime. Subclades C1 and C2 are indicated with green and blue branches, respectively. Subgroup C2a is shaded pink. The distribution of root-to-tip distances versus isolation date of all terminal nodes in the time-scaled tree is shown in the inset graph.

We identified a roughly 40%/60% split between isolates belonging to subclades C1 versus C2. Due to the persistence of both subclades, we investigated if these subclades differed in infection sites and antimicrobial resistance (AMR) gene presence. The only differences we observed in isolate source between the two clades, however, were slightly more blood isolates belonging to subclade C2 and slightly more respiratory isolates belonging to subclade C1 (Table S1). We identified acquired AMR genes in all genomes in our dataset, and then examined the AMR gene content in subclade C1 versus C2 genomes (Table S2, Fig. S2). We found that subclade C1 isolate genomes encoded slightly more AMR genes compared to subclade C2 genomes, however the difference was not significant (mean 7.8 vs. 7.1 genes, *P*=0.178). We also observed differences in the prevalence of individual genes conferring resistance to several different antibiotic classes between the different subclades, including aminoglycosides, antifolates, macrolides, and sulfonamides (Fig. S2).

### Minimal gene enrichment in subclade C1 and C2 genomes

We performed a pan-genome analysis for the 138 genomes in clade C using Roary^13^ to identify genes that may be beneficial in clade persistence. Among the 11,587 genes in the clade C pangenome, 3,429 genes were shared among all clade C genomes, representing 70.3% of the average number of genes among genomes in this clade (Table S3). Using an 80%/20% enrichment cut-off, there were only 13 genes that were enriched among subclade C1 genomes (Table S4), and no genes were enriched among subclade C2 genomes, perhaps because this subclade was larger and more diverse than subclade C1. Nearly all the 13 genes enriched among subclade C1 genomes appeared to be plasmid-encoded and were predicted to encode hypothetical proteins (Table S4).

Within subclade C2, we identified 56 genes that were specific to the *fimH*5 allele-carrying subgroup we designated as C2a (Fig. S3, Table S5). These genes appeared to be associated with several transposable units carrying carbohydrate and lipid metabolism genes as well as cell wall and cell membrane biogenesis genes (Table S5). We also identified a group of 27 subclade C2 genomes isolated between 2007 and 2018 that resided on the same phylogenetic branch, clustered together by accessory gene content, and carried 182 group-specific genes that we designated subgroup C2b (Fig. S3, Table S6). Approximately one third of these genes were associated with prophages, and 32 genes were predicted to reside within transposons. The remaining genes with annotated functions included carbohydrate transport and metabolism genes, antibiotic and heavy metal resistance genes, toxin genes, and cell envelope-associated factors (Table S6).

### Convergent evolution in subclades C1 and C2

We analyzed core genome non-synonymous SNPs in non-recombined genes among all isolates in each subclade to identify genes with multiple, independent SNPs in different isolates (Fig. 3, Table S7, Table S8). We focused on genes that had at least three non-synonymous SNPs among subclade C1 genomes (Fig. 3A), and at least four non-synonymous SNPs among subclade C2 genomes (Fig. 3B), as these genes would be unlikely to accrue so many mutations due to chance alone. Among subclade C1 genomes, the hydroxyacylglutathione hydrolase gene *gloB* and the peptidoglycan D,D-transpeptidase gene *ftsI* both possessed three different non-synonymous SNPs in three different isolates, and the undecaprenyl-phosphate 4-deoxy-4-formamido-L-arabinose transferase gene *arnC* possessed four different non-synonymous SNPs in five different isolates (Fig. 3C, Table S7). Both *ftsI* and *arnC* contribute to cell wall assembly, while *gloB* is involved in methylglyoxal detoxification^14^. Among subclade C2 genomes, two genes encoding hypothetical proteins (*DVT980_3104* and *DVT980_4259*) each possessed four different non-synonymous SNPs (Fig. 3C). One of these proteins (DVT980_3104) was similar to the ribosome association toxin encoded by *ratA* and was mutated in four different isolates, while the other protein (DVT980_4259) was similar to the enterobactin siderophore exporter encoded by *entS* and was mutated in 19 isolates (Table S8). The peptidoglycan D,D-transpeptidase gene *ftsI* possessed five different non-synonymous SNPs in five different subclade C2 isolates, none of which overlapped with the three *ftsI* mutations detected in subclade C1 isolates. Two different mutations were detected at amino acid position 413 in *ftsI* (Ala413Val and Ala413Thr), strongly suggesting adaptive evolution of this gene. Finally, the yersiniabactin/pesticin outer membrane receptor gene *fyuA* possessed eight different non-synonymous SNPs in nine different C2 isolates; such a high number of independent mutations also suggests strong selection acting on this gene.

**Figure 3.**
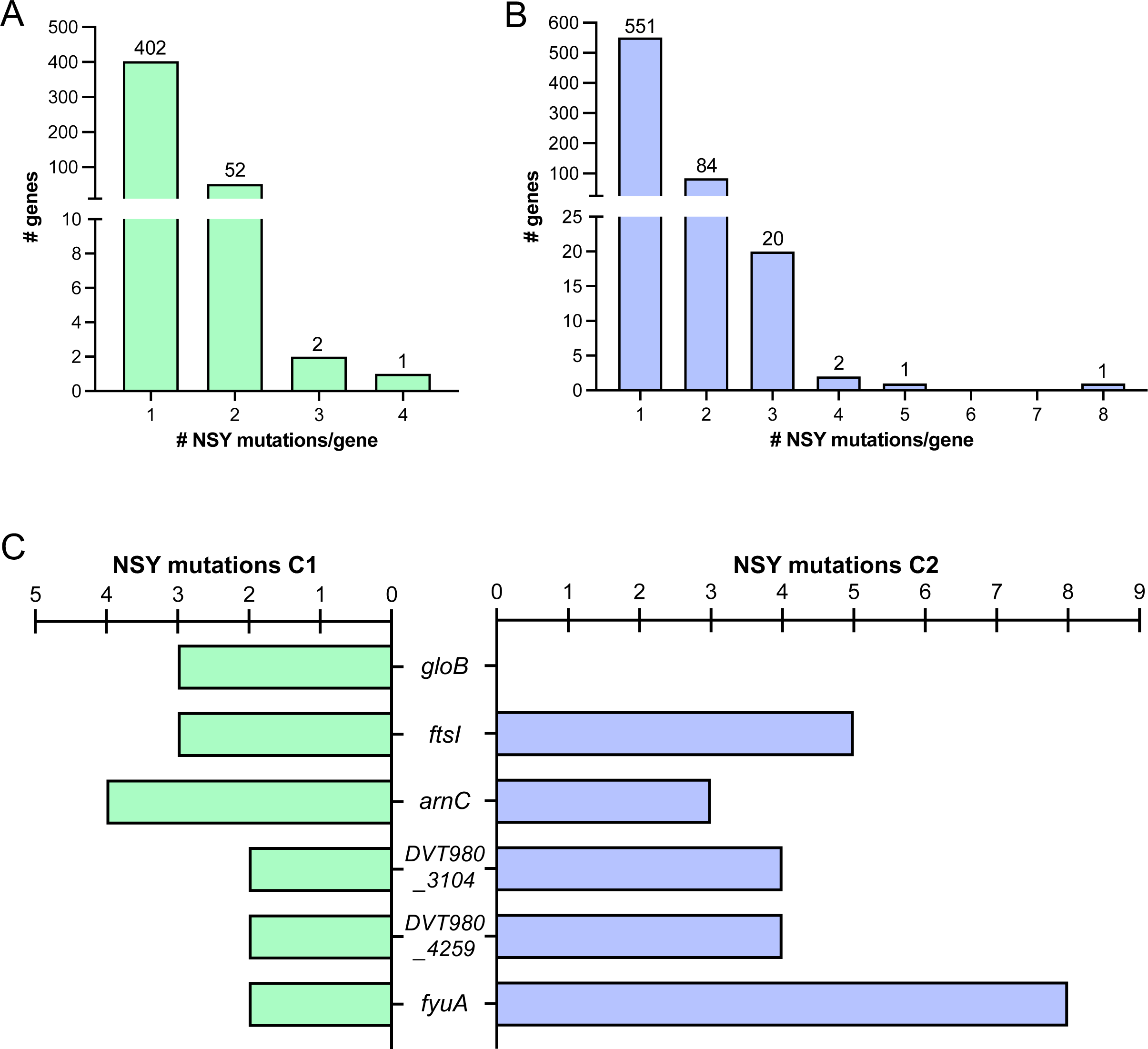
Genes putatively under selection among clade C *E. coli* isolates. Enrichment of nonsynonymous (NSY) mutations among subclade (A) C1 and (B) C2 genomes. Frequency distributions show the number of genes with one or more NSY mutation detected. (C) Genes with at least three unique NSY mutations in subclade C1 genomes or at least four unique NSY mutations in subclade C2 genomes. The number of different mutations detected in each gene among the genomes in each subclade is shown.

### ST131 clades carry diverse ESBL genes on both plasmids and the chromosome

To examine the diversity of ESBL genes carried by the isolates we collected, we performed BLASTP searches against the ResFinder database^15^. A total of twelve different ESBLs were detected, including CTX-M, SHV, and TEM family enzymes (Fig. 4A, Table S9). The most common ESBL enzyme detected was CTX-M-15, which was found in 93 genomes and was dominant in subclade C2 (79/83, 95.18%). Outside of subclade C2, CTX-M-15 was also found in nine subclade C1 genomes and in one clade A genome (Fig. 4A). The second most common ESBL enzyme detected was CTX-M-27, which was found in 32 genomes and was the most prevalent enzyme detected in subclade C1 (26/51, 50.98%) and clade A (6/12, 50%). CTX-M-27 was first detected in 2013, and was the dominant ESBL type identified in subclade C1 and in clade A in 2017 and 2018 (Table S1). The third most common enzyme we detected was CTX-M-14, which was found in nine genomes and was not associated with any specific clade or subclade (Fig. 4A). The remaining ESBL enzymes detected were CTX-M-2 (n=3), CTX-M-24 (n=3), CTX-M-1 (n=1), CTX-M-3 (n=1), SHV-12 (n=7), SHV-7 (n=1), TEM-19 (n=2), TEM-12 (n=2), and TEM-10 (n=1). One isolate (EC00670, belonging to subclade C2) was found to encode both CTX-M-14 and CTX-M-15 enzymes.

**Figure 4.**
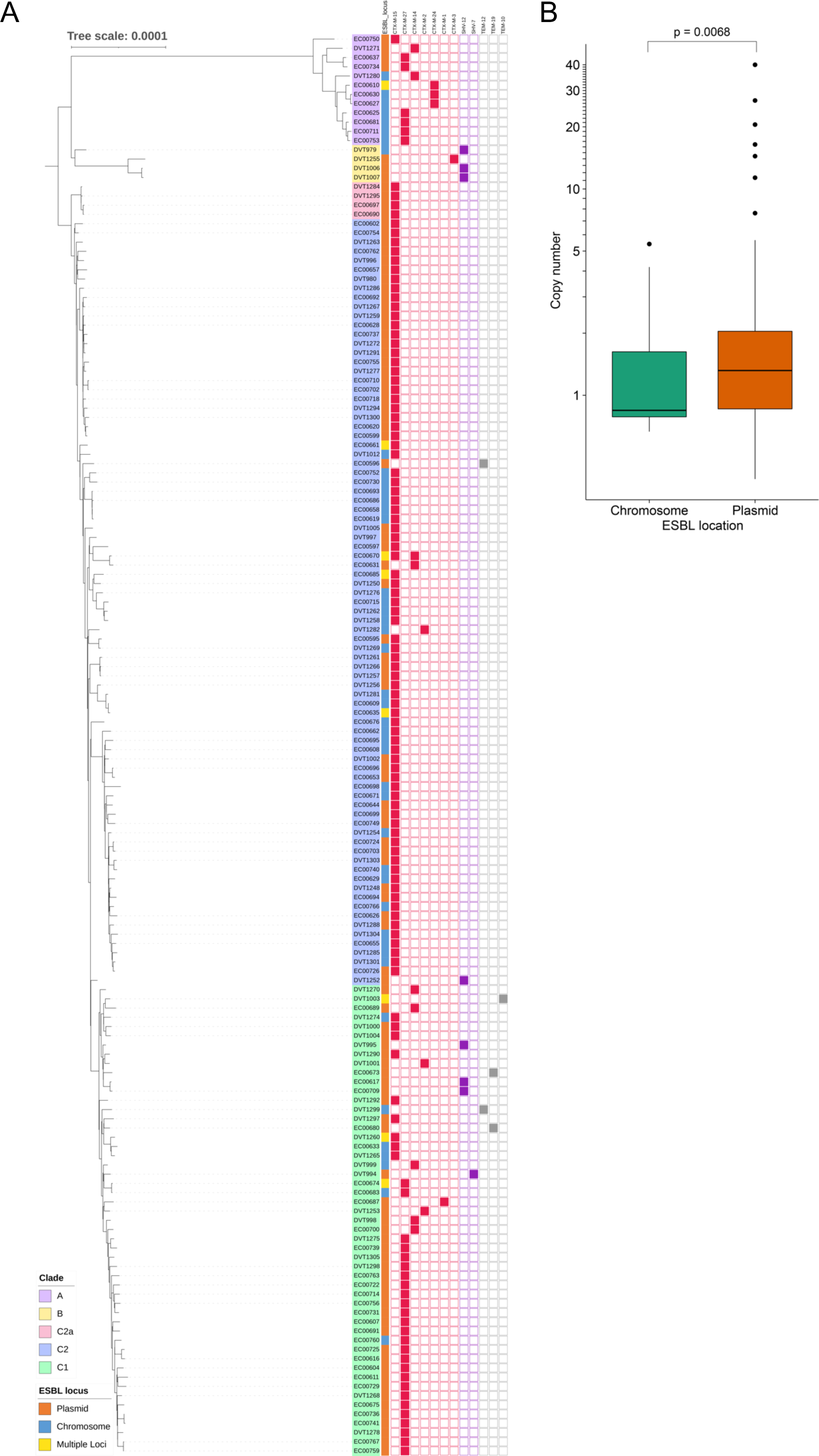
ESBL gene diversity, genomic location, and copy number. (A) Distribution of ESBL genes. ESBL locations (plasmid/chromosome/multiple loci) and types are shown as color blocks next to the isolate names. (B) Box plot showing ESBL gene copy number in isolates predicted to encode an ESBL gene on the chromosome or on a plasmid. *P*-value was calculated using a two-tailed t-test.

While ESBL genes are carried on MGEs, these elements can reside on plasmids or be integrated into the chromosome^1^. We assigned a putative genomic location of the ESBL enzyme in each isolate in our dataset using the MOB-RECON tool in MOB-Suite, which predicted whether ESBL-encoding contigs in each genome represented plasmid or chromosome sequences^16,17^. The majority of isolates (105/154, 68%) were predicted to carry ESBL genes on plasmids, while 46/154 (30%) were predicted to carry ESBL genes on the chromosome (Fig. 4A). The remaining isolates (3/154, 2%) were predicted to encode ESBL enzymes on both plasmids and the chromosome. Next, we used the 45 genomes that were hybrid assembled to examine the diversity and distribution of ESBL-encoding plasmids in our dataset. Among these 45 genomes we identified 35 ESBL-encoding plasmids, most of which belonged to the F family (Table S9). We then searched for each of these plasmids in all genomes in our dataset, and found that 11 plasmids were likely present in more than one isolate (Fig. S4). Four different *bla*_CTX-M-15_-carrying plasmids were found among subclade C2 genomes exclusively, while six of the other seven plasmids were found in isolates belonging to multiple clades. A total of 33 isolates that had ESBL enzymes predicted to be plasmid-encoded did not match to any of the 35 resolved ESBL-encoding plasmids using the identity and coverage cut-offs we employed (detailed further in the Methods), and likely contain different plasmid sequences.

Among the 45 hybrid assembled genomes, we identified eight genomes that had ESBL genes at more than one locus (Fig. 4A). The EC00610 genome carried three separate loci encoding CTX-M-24, all of which were on the chromosome. The EC00661 genome carried three loci encoding CTX-M-15, two of which were on chromosome and one of which was on a plasmid. The DVT1260 genome also carried two chromosomal loci encoding CTX-M-15, while the EC00685 and EC00635 genomes both encoded one CTX-M-15 locus on the chromosome and another locus on a plasmid. The EC00670 genome encoded one CTX-M-14 locus and one CTX-M-15 locus, each on two different plasmids, and the DVT1003 genome carried two loci encoding TEM-10 on two different plasmids. Finally, the EC00674 genome carried two loci encoding CTX-M-27 on the same plasmid.

To assess ESBL copy number variation in the isolates we collected, we quantified the estimated ESBL gene copy number in each genome by comparing Illumina sequencing read depth of the ESBL gene with the read depth of all single copy genes in the core genome (Table S10). We found that estimated ESBL copy numbers varied from 0.39x to 40x, with a median copy number of 1.15x. Isolates with chromosomal ESBL genes had an average ESBL copy number of 1.34x and a standard deviation of 1.06x, while isolates with plasmid-encoded ESBL genes had an average ESBL copy number of 2.73x and a standard deviation of 5.28x (Figure 4B). ESBL copy numbers were significantly higher among isolates with plasmid-encoded ESBLs (*P* = 0.0068).

### ESBLs are flanked by mobile elements that vary by enzyme type

To understand the genetic diversity of the elements carrying ESBL genes among the isolates we collected, we analyzed the genetic regions flanking the ESBL genes in each isolate in our study. Most assembled genomes allowed for examination of the genes immediately upstream and downstream of the ESBL enzyme (Fig. 5, Fig. S5). We found that *bla*_CTX-M-15_, which was present in 94% of subclade C2 isolates, very frequently resided in a conserved 3-kb region that was integrated into both plasmids and the chromosomes of different isolates (Fig. 5). We classified the *bla*_CTX-M-15_-flanking regions based on similarities in their gene organization and orientation, and identified four different MGE types. The first *bla*_CTX-M-15_-harboring MGE was found in isolates of clades A and C, and consisted of an IS*Ecp1* transposase and a small ORF with unknown function upstream of *bla*_CTX-M-15_ (Fig. 5A). This MGE was similar to the IS*Ecp1*-*bla*_CTX-M-15_-ORF477 transposition unit reported by Stoesser et al.^6^. The second MGE included the same upstream IS*Ecp1* transposase gene and small ORF with unknown function, as well as a Tn*2* transposase gene downstream of *bla*_CTX-M-15_ (Fig. 5B). This MGE was similar to the putative *bla*_CTX-M-15_ source element (Tn*2*-IS*Ecp1*-*bla*_CTX-M-15_-ORF477-Tn*2*) reported by Stoesser et al.^6^. A third MGE was found exclusively on plasmids, and was flanked on either side by IS*26* elements (Fig. 5C). The fourth MGE was only present in subclade C2 genomes, and was found on predicted chromosomal contigs, however it appears to have integrated at different chromosomal positions in different isolates (Fig. 5D).

**Figure 5.**
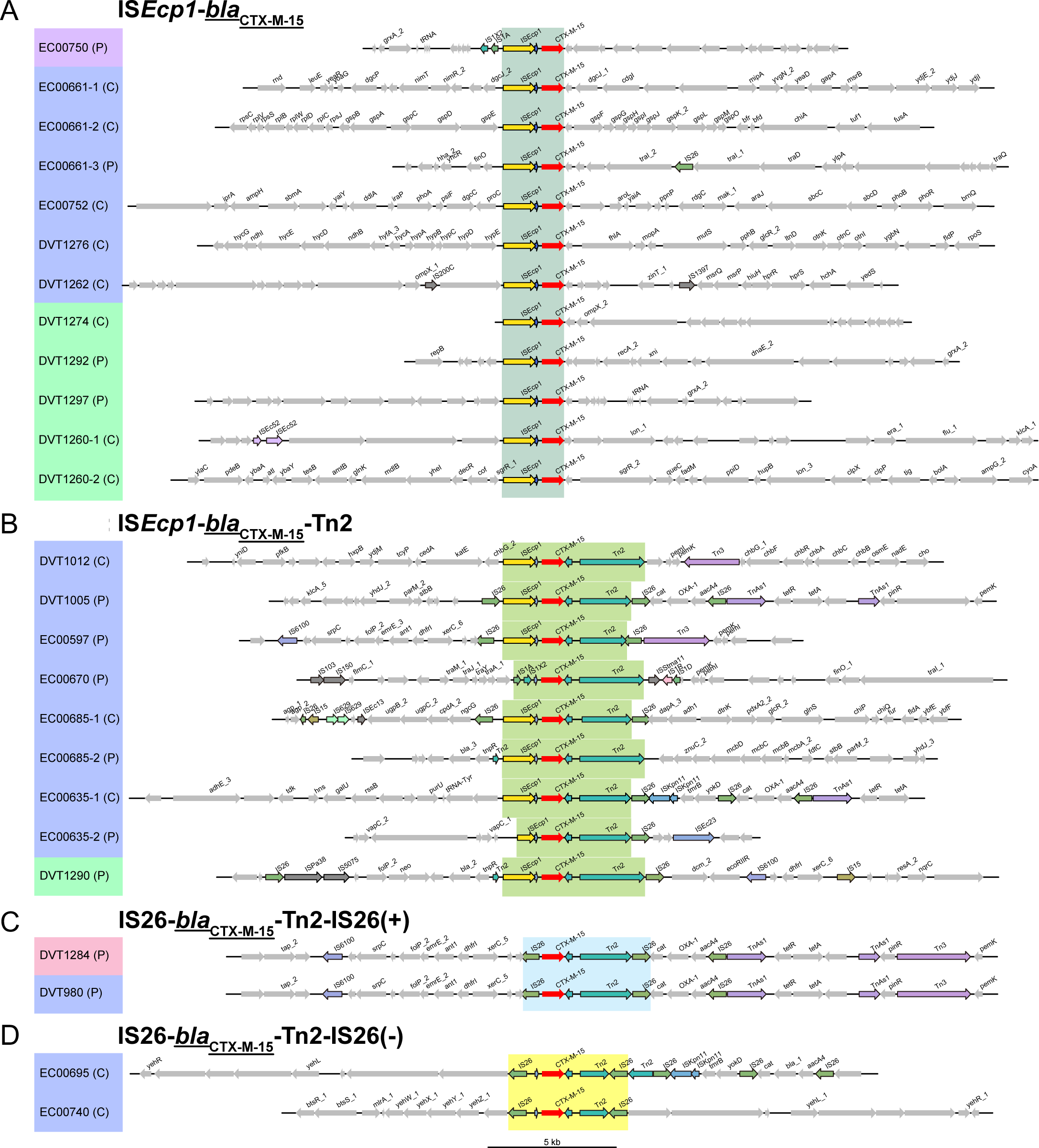
Regions flanking *bla*_CTX-M-15_ among ST131 *E. coli* isolates. (A-D) Genomic context of different *bla*_CTX-M-15_-carrying MGEs is shown. Isolate names are shaded based on their phylogenetic clade assignments (clade A=purple; subclade C2=blue; subclade C1=green; subgroup C2a=pink). The genomic location of each sequence is indicated (C=chromosome, P=plasmid) and *bla*_CTX-M-15_ genes are colored red. Genes were annotated with Prokka, and genes with predicted functions are labeled. Genes associated with MGEs and transposases are highlighted with black outlines, and are colored if found in more than one region. Regions that were used for MGE classification are shaded in each panel.

Apart from *bla*_CTX-M-15_, a variety of different MGEs were found to carry the other ESBL genes we detected (Fig. S5). *bla*_CTX-M-27_ was found on at least three different MGEs, and was associated with IS*15* and Tn*3* elements (Fig. S5A). Both *bla*_CTX-M-14_ and *bla*_CTX-M-24_ were found on the IS*Ecp1* MGE that also carried *bla*_CTX-M-15_ (Fig. S5B, S5C). Finally, *bla*_SHV-12_ was frequently found on a larger MGE that was flanked by IS*26* and contained additional carbohydrate metabolism genes (Fig. S5D).

## Discussion

In this 15-year study, we examined the genomic diversity and evolutionary dynamics of 154 ESBL-producing ST131 *E. coli* isolates from UPMC, a large healthcare system. Due to the multidrug resistance reported in ST131, numerous groups have characterized the clade structure of this pandemic lineage. Prior studies have suggested that clade C emerged around 1990^6,10,11^. Similarly, we identified the estimated root date to be midway through 1988. Our collection was dominated by isolates belonging to subclades C1 (*fim*H30-R) and C2 (*fim*H30-Rx). We identified the persistence of both clades at an approximate 40%/60% ratio, respectively. This finding suggests that these two subclades can coexist within the patient population that we sampled. We did not identify a significant difference in the number of AMR genes between the two clades, however, we did observe differences in the prevalence of individual genes conferring resistance to several different antibiotic classes. These data suggest that while subclade C1 and C2 isolates do not differ in their total AMR gene abundance, more subtle differences in the types of resistance genes they encode might contribute to their coexistence in the patient population that we sampled^18^.

We sought to further investigate why the C1 and C2 subclades have stably coexisted over the last 30 years. While our data suggest that subclades C1 and C2 do not harbor clade-specific gene signatures, within subclade C2 we identified two groups that were each enriched for genes with potentially useful functions. These enriched genes may contribute to ongoing adaptation of subclade C2 in the Pittsburgh area. In addition to subclade-specifying genes, we also investigated whether distinct genes might be under positive selection in subclade C1 versus C2 genomes. We identified missense variants in *gloB* were only detected in subclade C1 genomes, suggesting that perhaps mutating this gene was only beneficial in the subclade C1 genetic background. Multiple independent mutations in *ftsI* and *arnC* were detected in both subclades, and might affect bacterial susceptibility to other cell wall-targeting antibiotics like carbapenems^19^, or membrane-targeting antibiotics like colistin^20^, respectively. The *ratA*-like toxin and *entS* siderophore exporter genes were also independently mutated in multiple isolates across both subclades. These mutations might serve to decrease bacterial virulence, which frequently occurs during chronic infection and host adaptation^21^. Lastly, mutations in *fyuA* were also detected in both subclades, however they were heavily biased toward subclade C2 genomes. Prior studies have shown that *fyuA* function is critical for biofilm formation in iron-poor environments like the urinary tract^22^; mutations that alter or abrogate *fyuA* function would be predicted to decrease iron scavenging and biofilm formation. Future studies of the functional consequences of *fyuA* mutations on bacterial virulence and host-pathogen interactions may produce additional insights as to why these mutations appear to be under selection in ESBL-producing ST131 *E. coli* from our setting.

In agreement with previous reports, we identified a strong association CTX-M-15 and subclade C2 and CTX-M-27 and subclade C1^4,18,23^. The first isolate harboring CTX-M-27 in our collection was identified in 2013, coinciding with the recent emergence of CTX-M-27 documented in Europe and Asia^5,24,25^. When we predicted the location of the 154 ESBL-positive isolates, roughly a third were identified on the chromosome. While only one prior study has reported the chromosomal integration of CTX-M-14 in *E. coli* isolates from Mongolian wild birds^26^, this phenomenon has been described in a previous report of *Klebsiella pneumoniae* blood isolates, where nearly a quarter showed chromosomally-encoded EBSLs^27^. This finding suggests that the integration of the ESBL enzyme onto the chromosomal might enhance stable propagation and expression.

In addition to carrying a wide variety of ESBL genes, the ST131 *E. coli* isolates we sampled also carry a large diversity of ESBL-encoding plasmids. Some of these were specific to individual ST131 subclades, while others were identified widely throughout the lineage. We identified instances where isolates carried multiple ESBLs, either on different plasmids and/or integrated onto the chromosome. These data suggest that ESBL enzymes are frequently present at multiple loci within ST131 genomes, however these features can be difficult to resolve from Illumina draft genomes. Given that nearly 20% of our hybrid assembled genomes encoded ESBL enzymes at more than one locus, it is very likely that there are additional isolates in our dataset that also encode ESBL genes at multiple loci. The significance of this is unclear but could be due to gene dosage, plasmid instability, and/or shifting selective pressures during infection and antibiotic treatment^28,29^.

Prior studies have demonstrated that copy number variation of antibiotic resistance genes like β-lactamases impacts antibiotic susceptibility and facilitates the evolution of antibiotic resistance^30,31^. Our findings of variable ESBL copy numbers among the isolates we sequenced suggests that antibiotic selection might have increased the ESBL-encoding plasmid copy number in some isolates. Alternately, plasmid instability or fitness costs could have decreased copy numbers in other isolates. These findings relate to gene abundance and not transcript or protein abundance, nonetheless we find that ESBL gene copy numbers were both higher and more variable in isolates with plasmid encoded ESBLs.

ESBLs in ST131 *E. coli* are most often carried by MGEs that are integrated into plasmids or the chromosome^32,33^. Similar to prior work, we have identified the regions flanking *bla*_CTX-M-15_ have been found to be well conserved, even in distantly related genomes^6^. Further, through characterizing a variety of different MGE with ESBLs, our findings indicate that ESBL genes in the isolates from our medical center are likely shuttled between bacteria by MGEs that vary by enzyme type. Additionally, these elements appear to have integrated at different locations on both the plasmid and chromosome. It is notable that we observed a wide variety of different MGEs among the ST131 ESBL-producing *E. coli* sampled from a single geographic location. This suggests that as in other locations^34,35^, no single ESBL enzyme or MGE type was dominant at our center during the study period.

## Conclusions

This study describes ongoing adaptation of the ST131 *E. coli* population sampled from clinical cultures of patients in a single hospital in Pittsburgh. While the vast majority of isolates we collected belonged to ST131 clade C, both subclades C1 and C2 appear to be stably maintained over time in our facility. Despite this stable maintenance, we found an abundant diversity of ESBL enzyme types and a vast array of different mobile elements carrying these enzymes on both plasmids and the chromosome. The diversity of antimicrobial resistance genes, movement of plasmids and other MGEs, and signals of adaptation we identified will be the focus of our future work in this area.

## Methods

### Sample collection

Clinical bacterial isolates were collected from patients at the University of Pittsburgh Medical Center (UPMC), an adult tertiary care hospital with over 750 beds, 150 critical care unit beds, more than 32,000 yearly inpatient admissions, and over 400 solid organ transplants per year. Bacterial isolates included in this study were collected from patients as part of routine clinical care and were collected before they otherwise would have been discarded. The study was designated by the University of Pittsburgh institutional review board as being exempt from informed consent. Isolates were collected from 2004 through 2018, and were identified as *E. coli* initially by the clinical microbiology laboratory. ST131 isolates were identified with PCR using lineage-specific primers on isolates collected between 2004 and 2016^8^, or through analysis of whole genome sequences generated by the Enhanced Detection System for Healthcare-Associated Transmission (EDS-HAT) project in 2016-2018^9^. Collection of bacterial isolates was approved by the University of Pittsburgh institutional review board. ESBL phenotypes were inferred by the presence of an intact β-lactamase enzyme predicted to have ESBL activity within the genome of each isolate. Single bacterial colonies were isolated, and were grown on blood agar plates or in Lysogeny Broth (LB) media prior to genomic DNA extraction.

### Whole-genome sequencing

Genomic DNA was extracted from each isolate using a Qiagen DNeasy Tissue Kit according to the manufacturer’s instructions (Qiagen, Germantown, MD). Illumina library construction and sequencing were conducted using an Illumina Nextera DNA Sample Prep Kit with 150-bp paired-end reads, and libraries were sequenced on the NextSeq 550 sequencing platform (Illumina, San Diego, CA) at the Microbial Genome Sequencing Center (MiGS). A total of 45 isolates were also sequenced on a MinION device (Oxford Nanopore Technologies, Oxford, United Kingdom). Long-read sequencing libraries were prepared and multiplexed using a rapid multiplex barcoding kit (catalog SQK-RBK004) and were sequenced on R9.4.1 flow cells. Base-calling on raw reads was performed using Albacore v2.3.3 or Guppy v2.3.1 (Oxford Nanopore Technologies, Oxford, UK).

Short and long reads (or short reads alone) were used as inputs for Unicycler to generate draft genomes^36^. Plasmid and chromosomal contigs were predicted with the MOB-RECON tool in MOB-Suite v3.1.7^16,17^, and Prokka 1.14.5 was used for genome annotation^37^. Illumina raw reads for all isolates have been submitted to NCBI under BioProjects PRJNA475751 and PRJNA874473. Hybrid assembled genomes have been submitted to GenBank with accession numbers listed in Table S1.

### MLST, *fimH*, *gyrA/parC*, and clade C2 SNP Genotyping

Multi-locus sequence typing (MLST) was performed with SRST2^38^. Typing of the *fimH* locus was performed by running BLASTN against the *fimH* sequence database downloaded from FimTyper^39,40^. To detect quinolone resistance-determining region (QRDR) mutations, amino acid residues 81-87 of *gyrA* and the 78-84 of *parC* were extracted and compared^41^. To detect clade C2-specific single nucleotide polymorphisms (SNPs), targeted regions of primer sets described previously^4^ were extracted from all genomes and were compared with BLASTN.

### Phylogenetic trees and the time-scaled phylogeny

Among hybrid assembled genomes, the earliest collected isolate (DVT980) was used as a reference genome for Snippy v 4.6.0 to identify SNPs among the isolates using short read data and to generate a core SNP alignment (https://github.com/tseemann/snippy). The alignments were used as input for RAxMLHPC v 8.2.12 with [-m ASC_GTRCAT --asc-corr=lewis -V] flags to generate phylogenetic trees^42^. ClonalFrameML v1.12 was then used to filter recombinogenic regions^43^. Resulting trees were visualized with iTOL v6.3^44^ or FigTree v1.4.4 (https://github.com/rambaut/figtree/). Branch bootstraps supporting the clade C phylogeny were evaluated using RaxMLHPC with 100 rapid bootstrapping replicates with [-m ASC_GTRCAT -f a --asc-corr lewis -V] flags. Estimation of evolutionary rate and a time scaled phylogeny of clade C isolates was generated with TreeTime v0.9.2^12^, using a phylogenetic tree, ClonalFrameML-trimmed alignment, and the collection dates of the 138 isolates in clade C as input.

### ESBL gene detection and copy number variation

Amino acid sequences of all protein coding genes annotated by Prokka were used as queries to run BLASTP against the ResFinder amino acid database^15,40^. Hits with 100% identity and 100% length coverage were then filtered and manually curated to only include ESBL genes. Isolates with less than perfect matches to a database entry were compared with the NCBI non-redundant protein sequences (nr) database with BLASTP. All ESBL enzymes reported are perfect protein sequence matches. To estimate the copy number of the ESBL gene(s) in each genome, Illumina raw reads were mapped to the assembled draft genome using BWA with default parameters^45^. The read depth covering each gene was then calculated via the MULTICOV function of BEDTOOLS v2.30.0, with the input BAM file generated by BWA and the BED file that includes all protein coding genes, tRNAs, and rRNAs^46^. To normalize read coverage, we used an AWK pipeline to calculate the reads per kilobase per million mapped reads (RPKM) for each gene based on the depth list output of BEDTOOLS. A list of single copy genes shared by all genomes included in this study was extracted from the <gene_presence_absence.csv> output file of Roary v3.13.0^13^. For each genome, the median RPKM value of the single copy genes was calculated using the median() function in R. ESBL gene copy number in each genome was estimated by dividing the RPKM value of the ESBL gene(s) by the median RPKM value of single copy genes for the same genome.

### ESBL-encoding plasmid detection and analysis of flanking regions

A list of ESBL-encoding reference plasmids was first generated from all hybrid assembled genomes and plasmid contigs identified by MOB-RECON v3.1.7^16,17^. Contigs predicted to be circular by Unicycler v0.5.0 but not recognized as plasmids were not included in the reference plasmid list. To reduce redundancy, plasmids sharing >95% nucleotide similarity (defined as the product of query coverage and nucleotide identity) and encoding the same ESBL gene were combined and only the longest plasmid was retained. The remaining reference plasmids were then queried in all genomes using BLASTN and hits that had >95% nucleotide similarity were retained. Results were then manually curated to remove hits in genomes predicted to encode ESBLs on the chromosome only and hits to reference plasmids harboring a different ESBL. Among Illumina-only genomes, if there were hits to multiple reference plasmids with the same ESBL, only the longest reference plasmid was reported. To assess ESBL flanking regions, DNA segments containing up to 15 genes upstream and downstream of each ESBL gene were visualized via the R package genoPlotR, and were manually aligned centering on the ESBL gene to visualize conservation and enable classification of ESBL-containing MGEs^47^.

### Identification of subclade-specific genes and SNPs for clade C

The 138 annotated genomes belonging to clade C, including four genomes in clade C2a, were used for pangenome analysis. The pangenome analysis tool ROARY was used to generate a gene presence and absence matrix (gene_presence_absence.csv). Genes enriched in each clade were identified as those that were present in more than 80% of isolates within the clade and less than 20% of isolates outside the clade. The pangenome matrix was visualized using the heatmap() function in R. Genes associated with prophages and transposons were identified using PHASTER and MobileElementFinder, respectively^48-50^. Snippy was used to identify SNPs among clade C1 and C2 isolates using the DVT980 (earliest collected isolate) hybrid assembled genome as a reference. SNPs found in genomic regions identified by ClonalFrameML as putative recombinations were then masked. SNPs located in clade C core genes were annotated with gene description and locus tag of the reference genome. SNPs were then examined manually to identify genes with repeated and independent mutations within each subclade.

## Acknowledgements

We gratefully acknowledge Jane Marsh, Akansha Pradhan, and Alecia Rokes for their helpful input throughout the course of this study. This work was supported by grant R01AI127472 from the National Institutes of Health (L.H.H.), grant DAA3-19-65600-1 from the US Civilian Research & Development Foundation (D.V.T), and by the Department of Medicine at the University of Pittsburgh, School of Medicine (D.V.T.). The funders had no role in study design, data collection and analysis, decision to publish, or preparation of the manuscript.

## Author Contributions

SC and DVT designed the study. LHH and YD provided bacterial isolates. SC, MPG, HRN, and CLM performed experiments and generated results. SC, EGM, and DVT wrote the manuscript. All authors reviewed the manuscript and approved of its contents.

---

**Figure S1.**
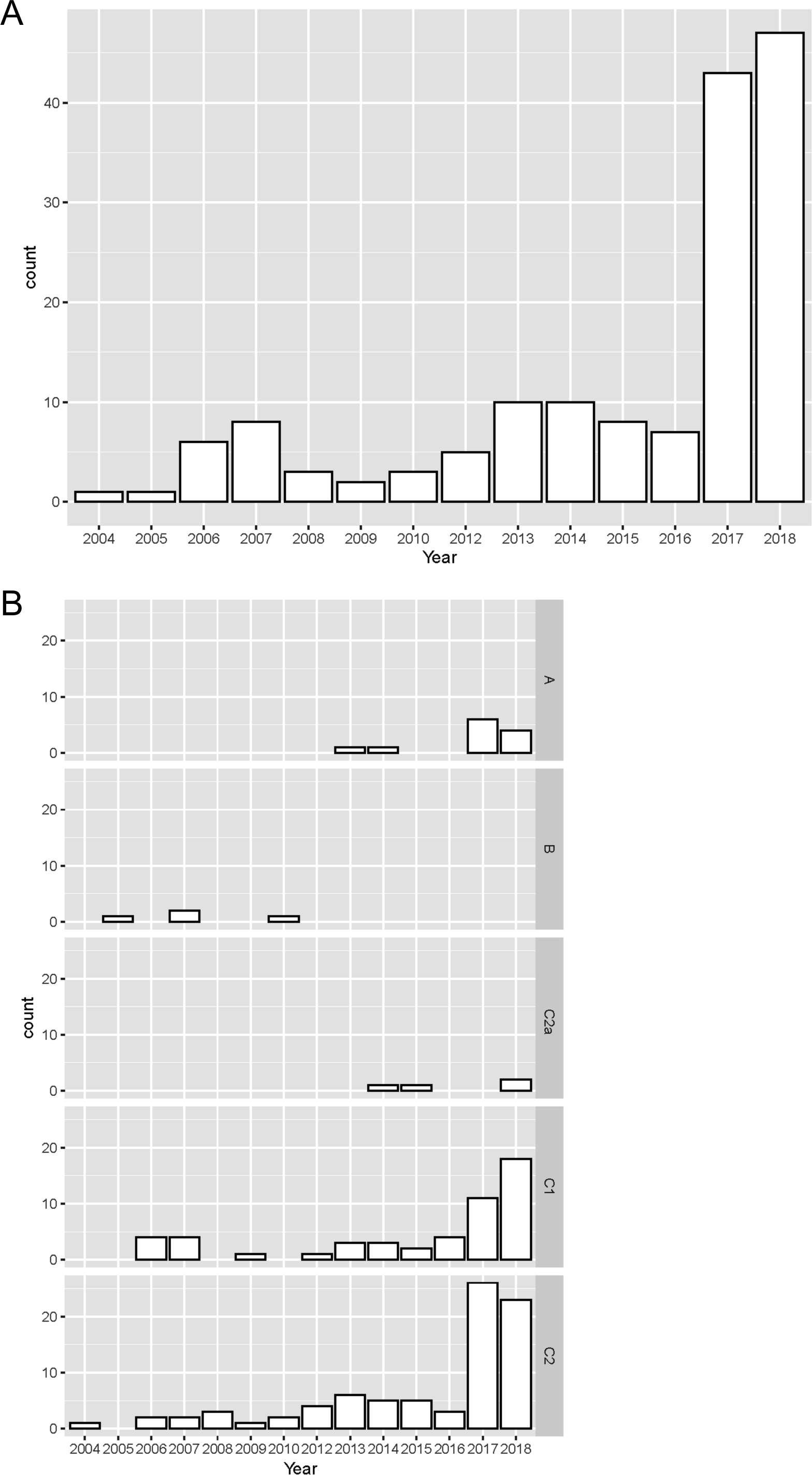
Timelines of ST131 isolate collection. (A) Total number of isolates collected each year. (B) Collection timelines for isolates belonging to each clade, subclade, and subgroup in the dataset.

**Figure S2.**
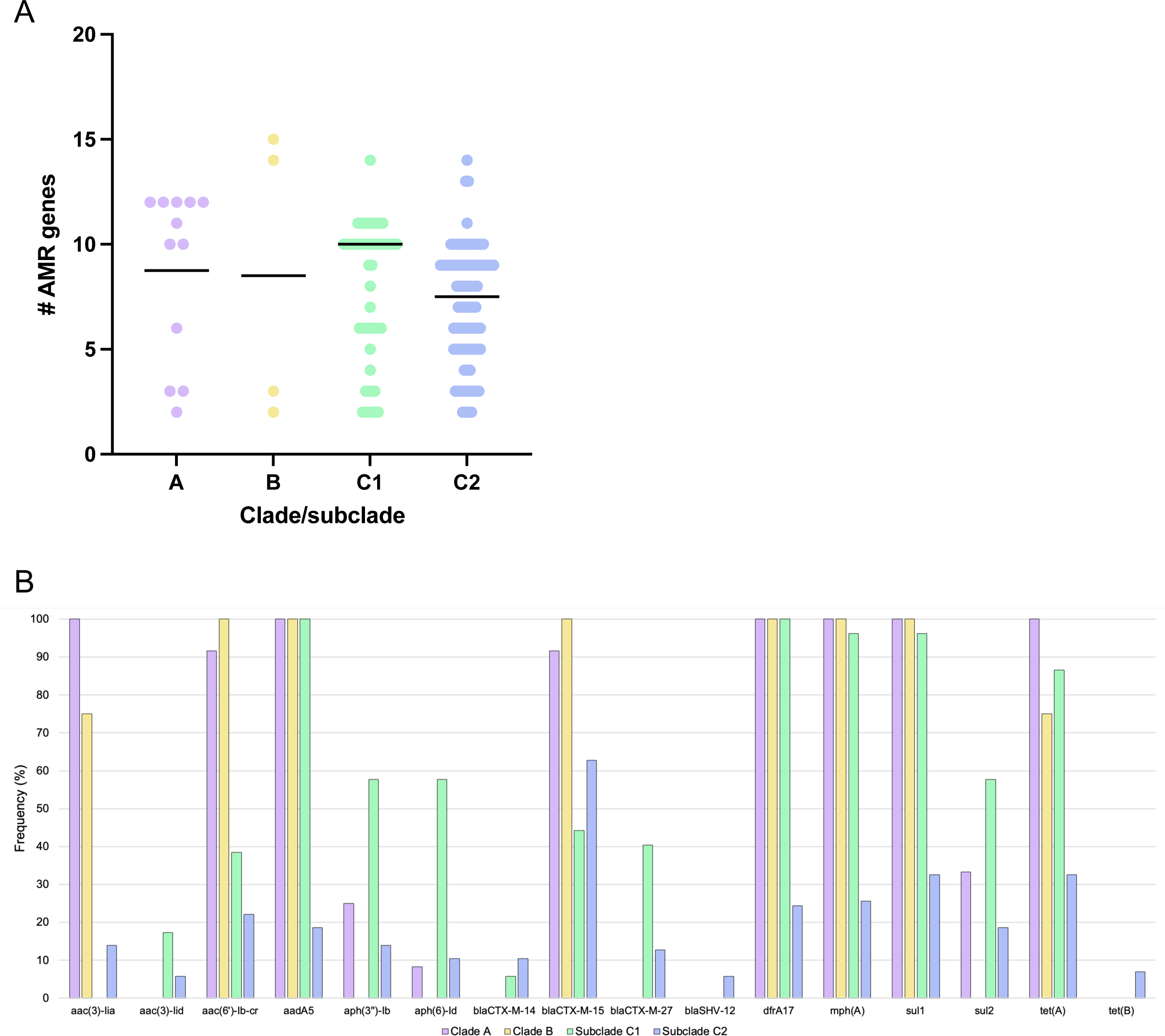
Differences in antibiotic resistance gene content between ST131 clades and subclades. (A) Antimicrobial resistance (AMR) gene abundance in isolates belonging to different ST131 clades and subclades. Horizontal lines show median values or AMR genes per genome in each isolate. AMR genes were identified by BLASTN to the ResFinder daabase. (B) Frequency of individual AMR genes among isolates in each clade or subclade. Genes with notable frequency differences between groups are shown. Complete data on AMR genes is provided in Table S2.

**Figure S3.**
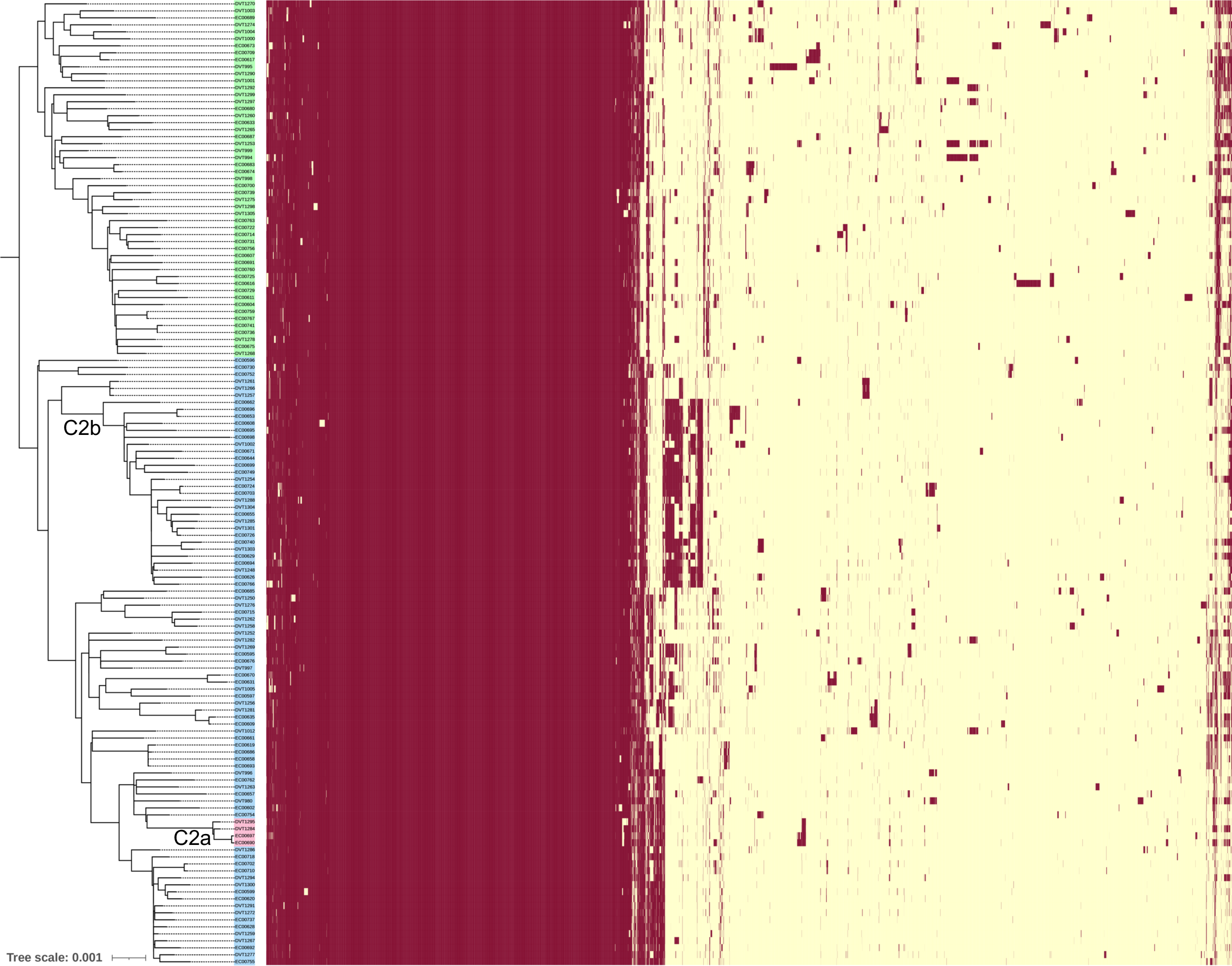
Pangenome analysis of 138 clade C ST131 *E. coli* isolates. Phylogenetic tree on the left was generated with RAxML using a core genome, post-ClonalFrameML SNP alignment. The tree was midpoint rooted to separate subclades C1 (green shaded) and C2 (blue shaded). The heatmap on the right shows the pangenome matrix generated by Roary. Each column represents one gene group, and each row represent one genome. The presence or absence of a gene in a given genome is shown as red or yellow, respectively. Subgroups C2a and C2b are labeled below the corresponding branches on the phylogenetic tree.

**Figure S4.**
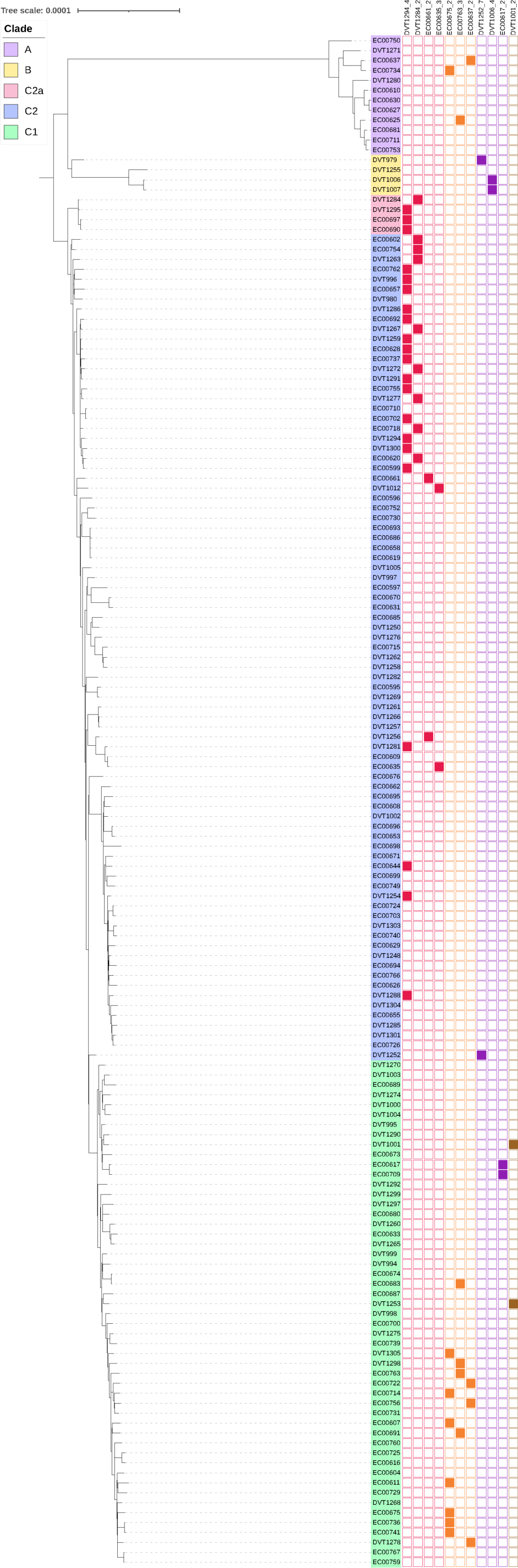
Distribution of ESBL-encoding plasmids among ST131 *E. coli* isolates. The core genome phylogeny is annotated with the presence of eleven ESBL-encoding reference plasmids that were detected in more than one genome in the dataset. Plasmids DVT1294_4, DVT1284_2, EC00661_2, and EC00635_3 harbor *bla*_CTX-M-15_ (red); plasmids EC00675_2, EC00763_3, and EC00637_2 harbor *bla*_CTX-M-27_ (orange); plasmids DVT1252_7, DVT1006_4, and EC00617_2 harbor *bla*_SHV-12_ (purple); and plasmid DVT1001_2 harbors *bla*_CTX-M-2_ (brown).

**Figure S5.**
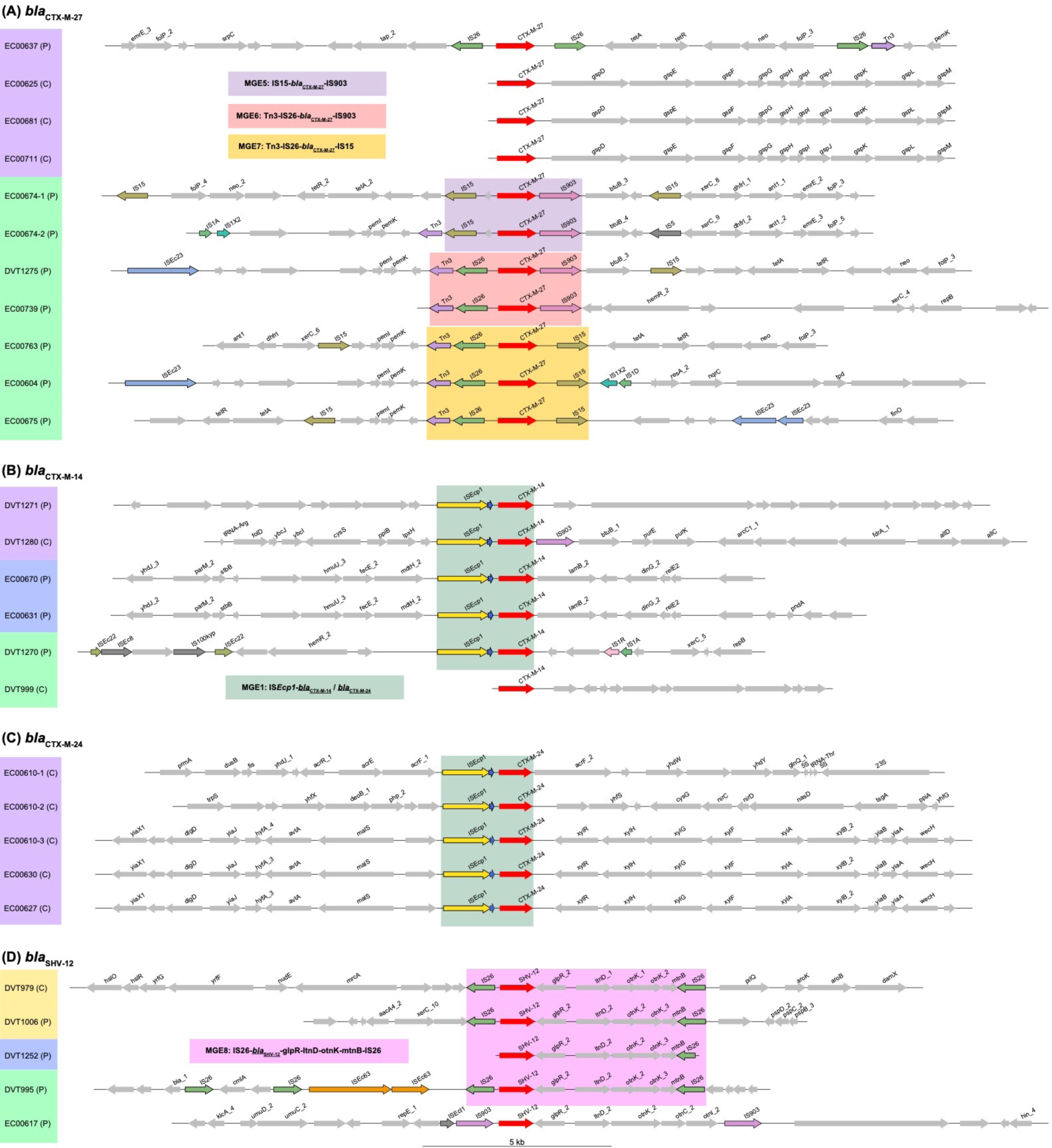
Regions flanking ESBL genes among ST131 *E. coli* isolates. (A-D) Genomic context of different ESBL-carrying MGEs is shown. Isolate names are shaded based on their phylogenetic clade assignments (clade A=purple; subclade C1=green; subclade C2=blue; clade B=yellow). The genomic context of each sequence is indicated (C=chromosome, P=plasmid) and ESBL genes are colored red. Genes were annotated with Prokka, and genes with predicted functions are labeled. Genes associated with MGEs and transposases are highlighted with black outlines, and are colored if found in more than one region. Regions that were used for MGE classification are shaded in each panel.

## Data Availability

Illumina raw reads and genome assemblies for all isolates have been submitted to NCBI under BioProjects PRJNA475751 and PRJNA874473. NCBI accession numbers for genome sequence data are listed in Table S1.

